# An Ensemble Learning and Slice Fusion Strategy for Three-Dimensional Nuclei Instance Segmentation

**DOI:** 10.1101/2022.04.28.489938

**Authors:** Liming Wu, Alain Chen, Paul Salama, Kenneth W. Dunn, Edward J. Delp

## Abstract

Automated microscopy image analysis is a fundamental step for digital pathology and computer aided diagnosis. Most existing deep learning methods typically require post-processing to achieve instance segmentation and are computationally expensive when directly used with 3D microscopy volumes. Supervised learning methods generally need large amounts of ground truth annotations for training whereas manually annotating ground truth masks is laborious especially for a 3D volume. To address these issues, we propose an ensemble learning and slice fusion strategy for 3D nuclei instance segmentation that we call Ensemble Mask R-CNN (EMR-CNN) which uses different object detectors to generate nuclei segmentation masks for each 2D slice of a volume and propose a 2D ensemble fusion and a 2D to 3D slice fusion to merge these 2D segmentation masks into a 3D segmentation mask. Our method does not need any ground truth annotations for training and can inference on any large size volumes. Our proposed method was tested on a variety of microscopy volumes collected from multiple regions of organ tissues. The execution time and robustness analyses show that our method is practical and effective.

## 1. Introduction

Optical microscopes have been widely used for imaging microscopic organisms and are important for the understanding of subcellular structures and disease diagnosis [1, 2]. The advances in digital fluorescence microscopy enabled multi-channel high resolution 3D imaging by using a diffraction-limited laser beam which can image deeper subcellular tissue structures [3, 4]. The 3D volumes generated by fluorescence microscopy need to be quantitatively analyzed to obtain useful information [5]. Manual assessment of large-scale microscopy volumes is laborious and time-consuming.

Deep learning-based methods have shown significant performance for computer vision tasks such as image classification, object localization and segmentation [6]. One of the major challenges in analyzing 3D microscopy volumes is to accurately delineate the boundary of individual nuclei having high intraclass variability and densely overlapping distributions [7, 8]. Many deep learning methods for image segmentation such as encoder-decoder-based networks typically require post-processing such as watershed or morphological operations to separate touching objects which may result in unstable results [9, 10]. Large computational resources such as large amounts of GPU memory are also necessary. One solution to reduce computational complexity is to process the volume as 2D slices and then merge the results to form a 3D nuclei segmentation. Robustly merging or fusing the 2D nuclei segmentations without compromising the overall accuracy remains challenging since the 3D nuclei information is not learned properly. Ensemble learning has been widely used to increase the overall robustness of segmentation approaches as well as increasing the segmentation accuracy by integrating the voting results from different networks [11]. However, supervised learning methods require large amounts of annotated training samples to achieve accurate results. Due to the lack of large amounts of ground truth data, quantitative analysis for some applications needs to be conducted without ground truth annotation for training supervised learning models [12].

In this paper, we describe an ensemble method for 3D nuclei segmentation, known as Ensemble Mask R-CNN (EMR-CNN), that is based on a collection of Mask R-CNN models with different network architectures for detecting and segmenting 3D nuclei in fluorescence microscopy volumes. We propose a weighted 2D mask fusion technique for aggregating 2D detection results from different Mask R-CNN networks to achieve more accurate and robust 2D results. We describe a 2D to 3D slice fusion method for merging segmentation results from 2D slices to a 3D volume using an unsupervised clustering method. By using ensemble learning, we demonstrate that our method achieves high nuclei detection accuracy compared to other methods we examined. In addition, we use Generative Adversarial Networks (GANs) to generate synthetic 3D microscopy volumes for training our EMR-CNN. Therefore our approach does not require any hand annotated ground truth for training which will be more useful when the hand annotated data is limited.

## 2. Related Work

Many methods have been reported in the literature for nuclei segmentation. They can generally be divided into five categories: threshold-based methods, clustering methods, energy-based methods, region-based methods, and machine learning methods [13].

The thresholding methods such as Otsu’s method try to determine a threshold that separates the foreground and background pixels by minimizing the intraclass intensity variance [14]. The typical region-based method is watershed [15], which treats the grayscale image as a topographic landscape with ridges and valleys, and the watershed transform is used to build barriers on the ridges to separate water source from different regions. The use of Otsu’s thresholding and watershed has been a popular combination for microscopy image segmentation [16, 17, 18]. Energy-based methods known as “Active contours” seek an equilibrium between the foreground object and background pixels by iteratively moving a deformable spline to minimize an energy function [19, 20].

Unsupervised learning techniques such as k-means, agglomerative hierarchical clustering (AHC), fuzzy C-means (FCM), and mean shift clustering have been used to split touching nuclei [21, 22, 23, 24]. These clustering methods explore the structure of nuclei and aggregate the pixels with similar features into different nuclei instances. The methods described above have been implemented and integrated as ImageJ plugins as well as other open source tools for quantitative microscopy image analysis [25, 17, 20, 26].

More recently, deep learning-based methods have shown promising results for cellular image analysis [27]. The encoder-decoder networks such as U-Net [28] and Seg-Net [29] have been used in microscopy image segmentation [30, 31, 32, 33], which demonstrate significant improved performance over classical image segmentation techniques. To segment different nuclei instances, post-processing such as watershed or morphological operation with proper parameter tuning is required [34]. To address this issue, the network has been modified to learn the centroid and boundary information of individual nucleus using a voting mechanism [35] or vector gradient map [36]. Similarly, [37] models the nuclei as ellipsoids and directly estimates the centroids and radii of the nuclei for instance segmentation. Alternatively, instead of directly segmenting an entire image, top-down approaches such as Region Proposal Networks (RPN) first identify the regions of interest (RoIs) and then segment the RoIs to obtain instance segmentations [38].

Ensemble learning techniques have been used to improve the overall detection performance by combining multiple diverse detectors, which can compensate for the errors generated by individual detectors [39]. An ensemble Mask-aided R-CNN described in [40] uses a graph clique voting method for improving the detection performance. In [41] an ensemble learning method based on CNNs and random forest (RF) for blood vessel segmentation is presented. Similarly, in [42] a transform modal ensemble learning for breast tumor segmentation is described. In addition, [43] described a cross-modality fusion and feature learning level ensemble learning for multimodal medical image segmentation and demonstrated the superiority of feature fusion over network output fusion such as voting. A weighted boxes fusion method was described for aggregating detected bounding boxes from different object detection models [44].

For segmenting 3D microscopy volumes, directly using 3D CNN networks can generally obtain more accurate results since the 3D information is utilized. However, 3D methods require 3D annotated ground truth for training, which is difficult to obtain in practice, especially for 3D microscopy volumes. To address these issues, many 2D to 3D methods have been introduced, which first perform the segmentation on the x-, y-, and z-direction of a volume and then fuse the results. In [45] 3D vector gradients are estimated by averaging 2D vector gradients from a modified 2D U-Net from three different directions of a volume, and followed by a clustering method to group the pixels to 3D masks. Similarly, [46] uses majority voting to combine 2D segmentation results obtained from SegNet into a 3D segmentation. However, accurately and robustly aggregating these 2D segmentation into 3D masks remains challenging.

To obtain satisfactory segmentation results, deep learning methods typically need large amounts training images with corresponding ground truth annotations. Manually delineating 3D nuclei contours or even 2D nuclei contours is laborious in a microscopy volume even for an expert. To address these issues, data augmentation methods including elastic deformation [47], random intensity correction, and spatial transformation are commonly used [48]. However, these methods require an adequate number of existing ground truth images. Learning-based data augmentation techniques such as generative adversarial networks (GANs) [49] can generate synthetic data without ground truth images. In [32] and [50], nuclei segmentation masks are generated by modeling nuclei as 3D ellipsoids and 3D non-ellipsoids using Bézier curves. In [31], a synthetic microscopy image generation method that is based on a modified CycleGAN [51], known as Sp-CycleGAN, uses the binary nuclei segmentation masks to generate corresponding synthetic microscopy images. In this paper we will use GANs to generate synthetic microscopy volumes for training our proposed method.

## 3. PROPOSED METHOD

In this section we describe the proposed Ensemble Mask R-CNN (EMR-CNN). We denote *I* as a 3D volume with dimension *X* × *Y* × *Z*. 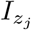 is the *j*-th slice of a volume on the *z* direction where *j* ∈ {1, …, *Z*}, and *I*_*z*_ denotes all 2D slices along the *z* direction of *I*. Also, *I*^orig^ denotes the original microscopy volumes. *I*^bi^ and *I*^label^ denote the binary segmentation masks and corresponding labels.

Let *E* be a collection or ensemble of 2D object detectors where *m*_*i*_ is a detector in *E* and *i* ∈ {1, …, *M*}. Given a 2D microscopy image 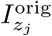 and an object detector *m*_*i*_, the detection results are denoted as 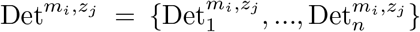 where 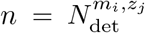 is the total number of detected objects in 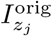 from *m*_*i*_, and each detection result 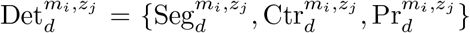 consists of a segmentation mask 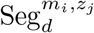, an object centroid 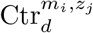 and a confidence score 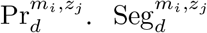 is a binary image of size *X* × *Y* where segmented pixels are highlighted with intensity 1. 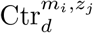 is a 2D coordinate indicating the centroid of the segmentation mask 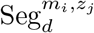, and 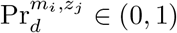 is the detection confidence score of the segmentation mask.

Also, let 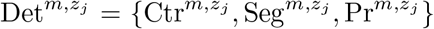 be the 2D detection results on 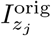 for all detectors in *E*. Our goal is to take all the detection results and fuse them together. Let 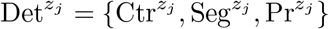 be the fused 2D detection result using ensemble 2D fusion method described in Section 3.3. Let Det^*z*^ represent a set of all 2D fused results for *I*^orig^, and Det = {Seg, Ctr, Pr} is the final 3D results fused from Det^*z*^. Next we take the fused 2D results Det^*z*^ and merge them to form our final 3D detection Det using a 2D to 3D slice fusion method described in Section 3.4.

### 3.1. Synthetic Data Generation

As we indicated earlier it is very difficult to obtain manually annotated microscopy volumes due to the tedious nature of the annotation process. We use a data augmentation process that consists of generating synthetic 3D microscopy volumes using GANs [51, 31]. As shown in Figure 1, the synthetic data generation module includes 3D ground truth nuclei segmentation masks generation and synthetic microscopy volume generation.

**Figure 1.**
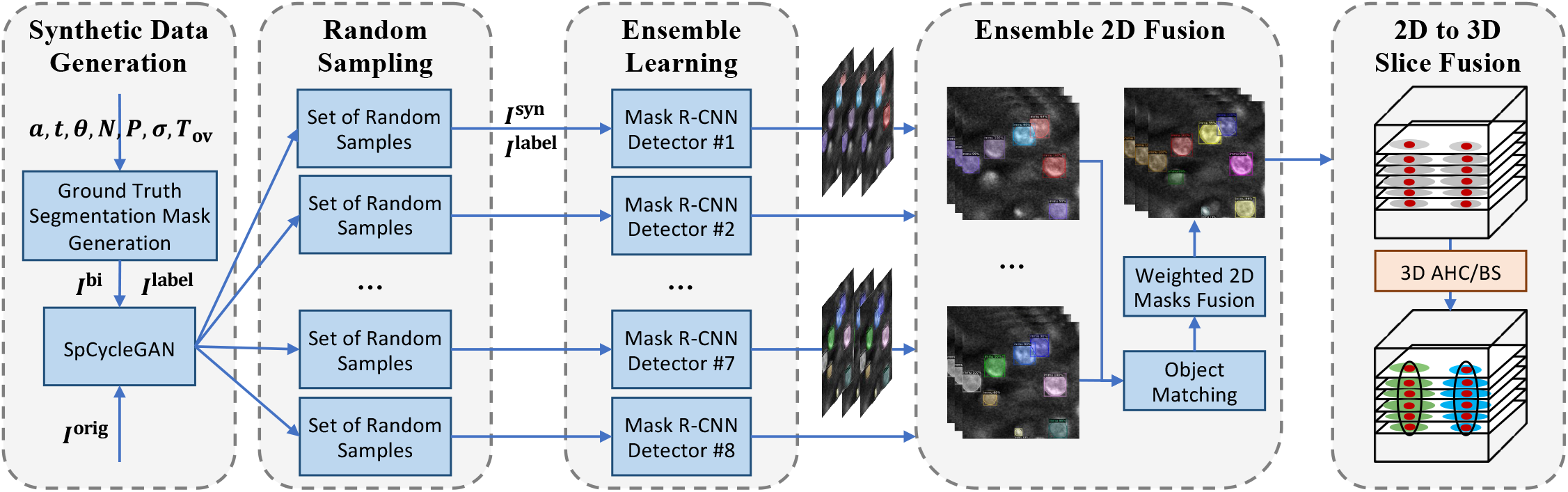
Overview of the proposed EMR-CNN for 3D nuclei instance segmentation using ensemble learning and slice fusion

#### Ground truth segmentation mask generation

We first generate synthetic 3D segmentation masks of nuclei that serve as the ground truth for the training data. Our approach is different from previous approaches described in [32, 50] in that we model each candidate nucleus as a deformed 3D ellipsoid parameterized by three parameters ***a, t***, and ***θ***, where ***a*** = (*a*_*x*_, *a*_*y*_, *a*_*z*_) defines three axis lengths of an ellipsoid, ***t*** = (*t*_*x*_, *t*_*y*_, *t*_*z*_) is the spatial translation that defines the location of a nucleus, and ***θ*** = (*θ*_*x*_, *θ*_*y*_, *θ*_*z*_) is the spatial rotation that defines the orientation of a nucleus. The parameters ***a, t*** and ***θ*** are randomly generated in a range shown in Table 1 for each candidate nucleus. These candidate nuclei are recursively added to an empty 3D volume *I*^*label*^ with an incremental unique intensity *k* that is used to distinguish different nuclei instances. The total number of nuclei is set to *N*, and the overlapping voxels of two nuclei must be less than *T*_ov_.

**Table 1.** Parameters for generating synthetic binary segmentation volumes. *r*_*x,y,z*_, *t*_*x,y,z*_, *θ*_*x,y,z*_ are randomly generated for each nucleus, and *P, σ, N* and *T*_ov_ are predefined values based on actual microscopy volumes

| Data | $r_{x,y,z}$ | $t_{x,y,z}$ | $\theta_{x,y,z}$ | $T_{\text{ov}}$ | $N$ | $P$ | $\sigma$ |
| --- | --- | --- | --- | --- | --- | --- | --- |
| $\mathcal{D}_1$ | (4, 8) | (1,128) | (0,2π) | 5 | 400 | 0 | 0 |
| $\mathcal{D}_2$ | (10, 14) | (1,128) | (0,2π) | 10 | 40 | 0 | 0 |
| $\mathcal{D}_3$ | (8, 10) | (1,128) | (0,2π) | 300 | 400 | 4 | 4 |

In the data we used for our experiments, we observed many nuclei are not strictly ellipsoids but more like “deformed” ellipsoids (Figure 2). To model these type of nuclei, we further deform the binary ellipsoids using an elastic transform [52]. Specifically, given a volume *I* of size *X* ×*Y* ×*Z* that needs to be deformed, we first generate a random coarse displacement, which is a matrix of size 3 × *P* × *P* × *P*, sampled from a normal distribution 𝒩(0, *σ*^2^). Then a displacement vector field *I*^vec^ was generated by interpolating the coarse displacement from size 3 × *P* × *P* × *P* to size 3 × *X* × *Y* × *Z* by cubic spline interpolation [53]. Then, *I*^vec^ is used to shift the voxels in *I* to obtain the deformed volume *I*^bi^ (Figure 2).

**Figure 2.**
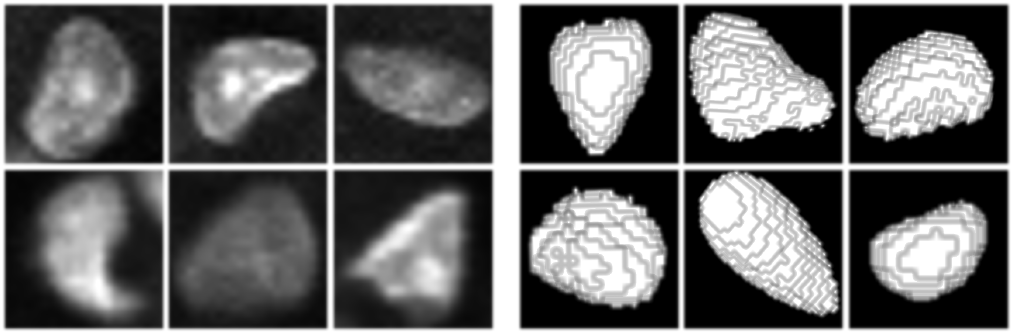
Real nuclei (left) and deformed ellipsoids generated using elastic transform (right)

#### Microscopy volume generation

We use the SpCycleGAN described in [31, 51] to generate the synthetic 3D microscopy volume. As shown in Figure 3, for SpCycleGAN training, we use all XY focal planes of unpaired original microscopy volumes *I*^orig^ and synthetic binary segmentation masks *I*^bi^. Once the SpCycleGAN is trained, we use *I*^bi^ as input to SpCycleGAN to generate a corresponding synthetic microscopy volume *I*^syn^. We generate the volume slice by slice and stack the results together.

**Figure 3.**
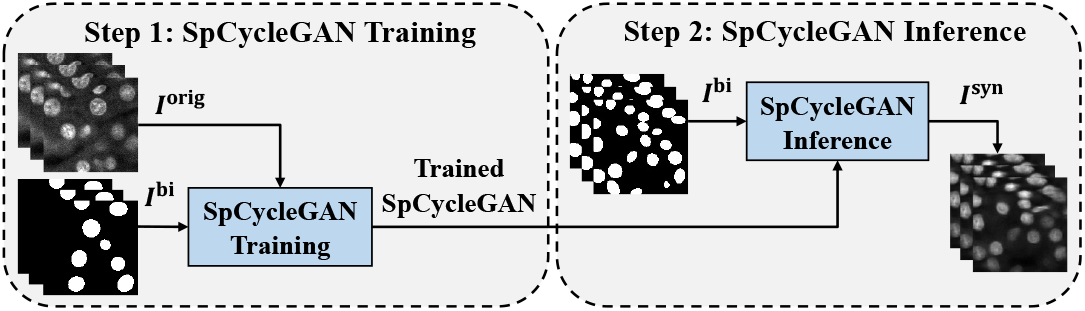
3D synthetic microscopy volume generation using SpCycleGAN

**Figure 4.**
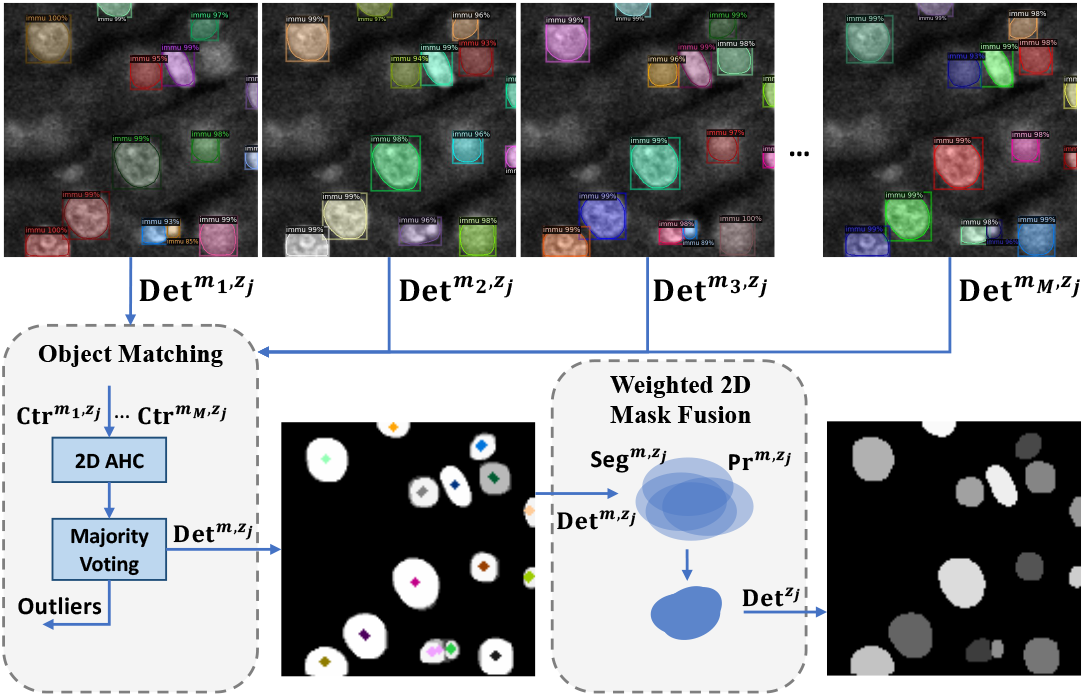
Overview of weighted 2D mask fusion

SpCycleGAN consists of 5 networks *G, F, H, D*_*A*_ and *D*_*B*_ where *G* is the generator that translates the image from binary domain to microscopy domain. Similarly, *F* translates the image from microscopy domain to the binary domain. *D*_*A*_ and *D*_*B*_ are two discriminators that are used to discriminate whether a given image is real or synthetic. The additional segmentation network *H* has the same architecture as *F* along with spatial constrained loss ℒ_sc_ were introduced in [31] to preserve the spatial shift of objects in the generated images. The entire objective loss function of SpCycleGAN is shown in Equation 1.

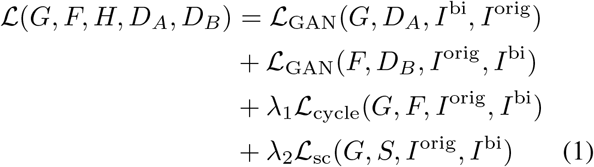

where ℒ_sc_ is the spatial constrained loss defined as a *L*_2_ norm shown in Equation 2.

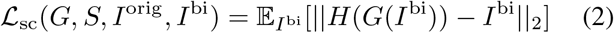

### 3.2. Ensemble Mask R-CNN: EMR-CNN

In order to increase the robustness and accuracy of nuclei segmentation, we propose a simple but effective method that trains a collection of *M* different but similar Mask R-CNN detectors implemented by [54]. The details of the training are described in Section 4.1. Our method includes ensemble 2D fusion that is able to fuse the 2D detection results from all detectors, and a 2D to 3D slice merging method that merges the detection results from fused 2D slices to 3D volumes.

### 3.3. Ensemble 2D Fusion

We propose a weighted 2D mask fusion method for fusing 2D detection results from different detectors into a final 2D detection. This is an extension of the weighted boxes fusion described in [55]. Instead of simply using Non-Maximum Suppression (NMS), we use the estimated confidence scores to compute the fused 2D segmentation mask which is represented by a probability map. A fused mask with new confidence score is generated. Given the detection results of a 2D slice 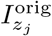 from *M* detectors, the goal is to fuse 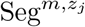 and 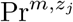into 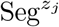 and 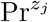.

#### Object matching

For an object, we need to identify all objects detected with the *M* detectors and then fuse the results. We use the agglomerative hierarchical clustering (AHC) with average linkage criterion to match the same segmented object in the segmentation masks 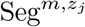. Specifically, we use Ward’s minimum variance implemented with Lance–Williams dissimilarity (L) [56] to determine which segmentation mask to be merged at each iteration. The Lance–Williams dissimilarity measures the similarity between an existing cluster and newly merged cluster. AHC will treat each sample as one cluster initially and merge the most similar sample pair based on the Lance–Williams dissimilarity until all samples are merged as one cluster. To evaluate the clustering performance, we define the mean intracluster distance *a*(*i*), and the mean intercluster distance *b*(*i*) for a given number of clusters *k* in Equation 3.

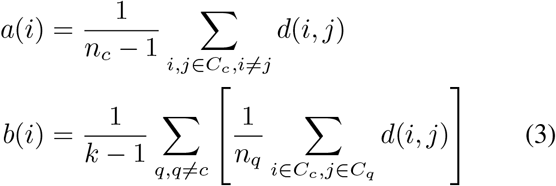

For a given segmentation mask centroid *i* and its cluster *C*_*c*_, *a*(*i*) measures the intracluster distance between *i* and other samples *j* within the same cluster. Similarly, *b*(*i*) measures the intercluster distance between *i* and other clusters *C*_*q*_. *n*_*c*_ is the number of elements in cluster *C*_*c*_. Note that *d*(*i, j*) is the Euclidean distance between two centroids. Finally, the Silhouette Coefficient (SC) is used to determine the ideal number of clusters 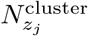. As shown in Equation 4, 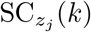 is the Silhouette Coefficient for the *j*-th slice given *k* as the number of clusters.

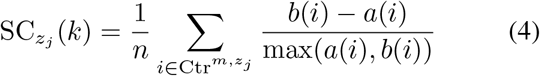

Our objective function can be formulated in Equation 5 to find the number of clusters *k* ∈ {*k*_min_, …, *k*_max_} with the highest Silhouette Coefficient.

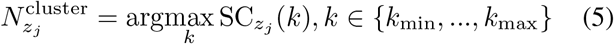

where *k*_min_ and *k*_max_ shown in Equation 6 are the minimum and maximum number of detection for all detectors and *σ*^2^ is the variance of the number detections for all *M* methods. Note that AHC requires there to be at least 2 potential clusters for unclustered centroids.

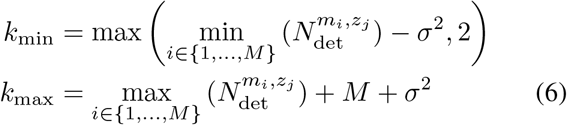

The majority voting mechanism is used to increase the robustness of the fusion by removing false positive detections or outliers. As shown in Equation 7, if the number of detections for an object in cluster *C*_*r*_ is less than *M/*2, the corresponding detection results will be removed.

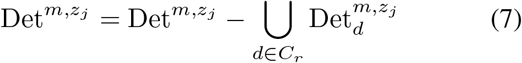

#### Weighted 2D mask fusion

To fuse the 2D segmentation masks from the *M* detectors, we propose weighted 2D mask fusion that takes the matched objects as input and outputs the fused 2D detections with corresponding confidence scores. This is shown in Equation 8.

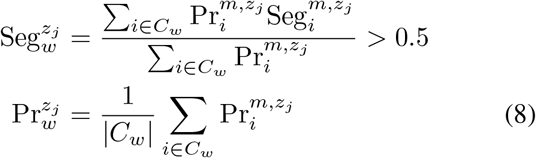

where *C*_*w*_ is the *w*-th cluster within which the detection results are matched. Then 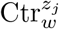 is the center of mass of 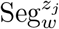.

### 3.4. 2D to 3D Slice Fusion

As described above, for a given an image slice 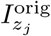 from the original volume, we obtain *M* 2D detentions and fuse them to form a 2D fused slice result 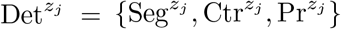. Our 2D to 3D slice fusion method merges 2D fused detections from all slices Det^*z*^ into a 3D detection Det = {Seg, Ctr, Pr}. As shown in Figure 1, we use two 2D to 3D slice fusion approaches, both based on the spatial location of the 2D centroid of the fused slices. The first approach known as blob-slice (BS) [57, 17] merges each 2D fused segmentation from the top slice to the bottom slice based on a predefined Euclidean distance of the centroid. The second approach uses agglomerative hierarchical clustering (AHC) described in Equation 3, 4, and 5 to cluster Ctr^*z*^. The confidence scores of the final 3D fused segmented objects are the average of confidence scores of corresponding 2D segmentations within the same cluster.

#### Slice merging

Since EMR-CNN operates on different slices of a volume independently without knowing the 3D nuclei structures in the *z*-direction, it may fail to detect the nuclei in a slice due to artifacts or the effect of point spread function. This results in a single segmented nucleus containing two or more disjoint connected components. We propose a technique for merging these disconnected components. For the 3D fused segmentation of a single nucleus Seg_*d*_, suppose a 2D fused segmentation 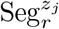 is missing on the *j*-th slice, and suppose the 2D fused segmentation for its neighbor slices are 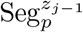 and 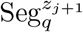, then the missing segmentations are given by the intersection of its two neighbor segmentations 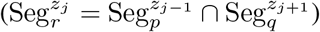.

## 4. EXPERIMENTAL RESULTS

### Datasets

In our experiments, we use three microscopy datasets denoted as 𝒟_1_, 𝒟_2_, and 𝒟_3_ for evaluation. The data is from various regions of a rat kidney using confocal fluorescent microscopy with fluorescence label (Hoechst 33342 stain). Original microscopy data were collected by Malgorzata Kamocka and Michael Ferkowicz at the Indiana Center for Biological Microscopy [58]. Microscopy 𝒟_1_ consists of one volume of *X* × *Y* × *Z* = 128 × 128 × 64 voxels, 𝒟_2_ consists of 16 volumes of size 128 × 128 × 32 voxels, and 𝒟_3_ consists of 4 volumes of size 128×128×40 voxels. These datasets were manually annotated using ITK-SNAP [59]. For training, we generate synthetic data for training EMR-CNN and other comparison methods. The synthetic 𝒟_1_, 𝒟_2_, and 𝒟_3_ are generated using 3 different trained SpCycleGANs (Figure 3). Each type of synthetic dataset consists of 50 volumes of size 128 × 128 × 128.

### 4.1 Experimental Setup

The training of SpCycleGAN requires unpaired 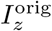 and 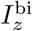 images. We first generate 54 binary segmentation masks *I*^bi^ for 𝒟_1_, 𝒟_2_, and 𝒟_3_ respectively, using the method described in Section 3.1. The axes lengths, translation distances, and rotation angles are randomly selected from uniform distributions having ranges *r*_*x,y,z*_, *t*_*x,y,z*_, and *θ*_*x,y,z*_, which are provided in Table 1, respectively. For synthetic microscopy volume generation, shown in Figure 3, we use all XY focal planes of 4 sub-volumes of *I*^orig^ along with 4 volumes of *I*^bi^ for training each SpCycleGAN model, respectively. We use the trained SpCycleGANs and the remaining 50 binary volumes to generate corresponding *I*^syn^ volumes for each dataset. Example images of the synthetic volumes are shown in Figure 5. The SpCycleGAN parameters are the same settings as described in [23, 31].

**Figure 5.**
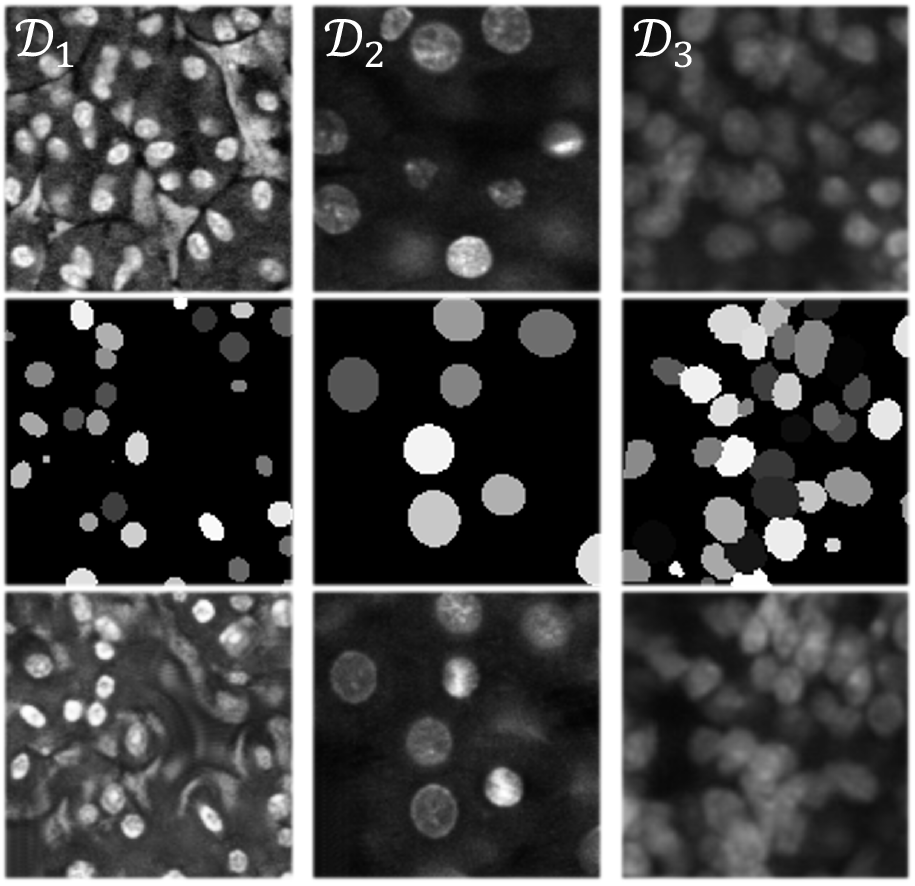
Original microscopy image (first row), synthetic nuclei segmentation mask (second row), and corresponding synthetic microscopy image (third row) for each dataset

### EMR-CNN training and inference

In our experiments, we trained 3 EMR-CNNs each consisting of *M* Mask R-CNN detectors that are initialized with different network architectures randomly chosen from ResNet-50, ResNet-101 [60], ResNeXt-101 [61], Feature Pyramid Network [62], and deep CNN layers [63]. All the networks are pretrained on the MSCOCO dataset [64] and are available at [54]. The networks are then retrained on a subset of the synthetic microscopy images and tested on actual microscopy volumes. The training subset is randomly chosen from the entire synthetic image training set. All ensemble models are trained on 4 TITAN XP GPUs in parallel with a base learning rate of 2.5*e*^−4^ and batch size of 2 for 2000 iterations. The remaining parameters, which are Mask R-CNN detector parameters, are set to their default values [54]. In addition, since the training images are of size 128×128 while the testing images can be arbitrarily large, directly inferencing on those large volumes may not generate accurate detection results (see Figure 6 left column) because the objects are downscaled. To address this, we propose a divide-and-conquer inference scheme that partitions the input volume to multiple 128×128×128 subvolumes with a 16-pixel border overlap (see Figure 6 right column). After the EMR-CNN inferencing stage, partial objects that lie on the overlapping boundaries of each partition are reconstructed based on their overlapping regions. Specifically, two objects on the partition boundaries that overlap by more than 10 pixels are merged into one object. Figure 6 middle column demonstrates how a naïve divide- and-conquer inferencing that does not utilize overlapping sub-volumes will result in segmentation errors at the border of the inference windows.

**Figure 6.**
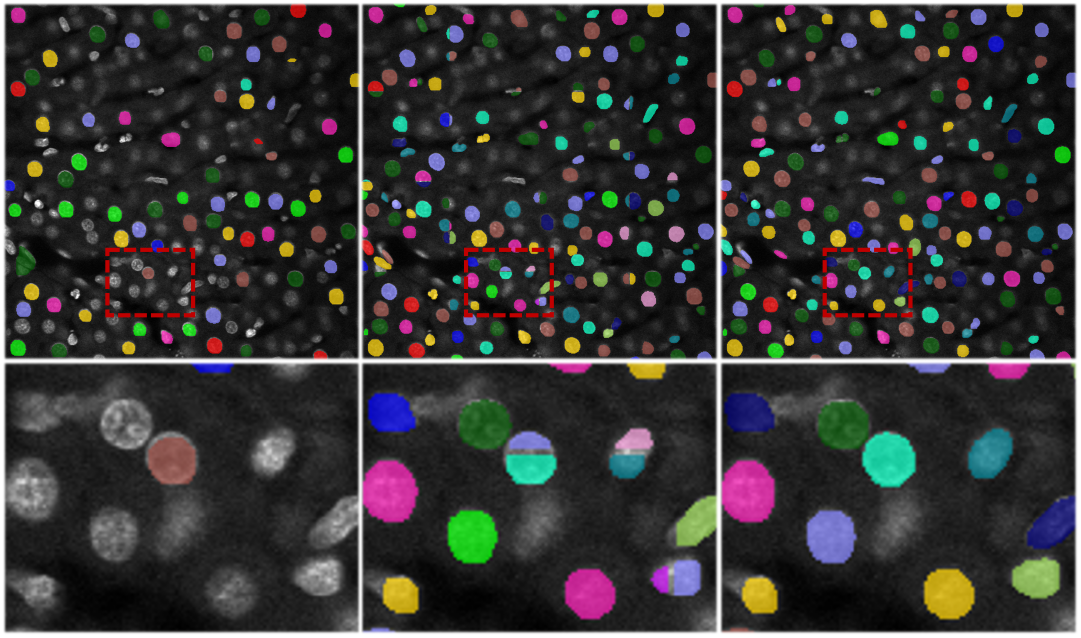
Segmentation results of EMR-CNN+AHC (M=4) on large microscopy 𝒟_2_. Left column: direct inference on each slice. Middle column: naïve segmentation based on non-overlapping partitions. Right column: results using our proposed divide-and-conquer inference method.

### 4.2. Quantitative Evaluation

To evaluate the accuracy of our method, we define that a ground truth nucleus is successfully detected if the Intersection-over-Union (IoU) between this ground truth nucleus and a detected nucleus is greater than an IoU threshold *t*. Then we count the True Positive (TP) detections (number of ground truth nuclei that are successfully detected), the False Positive (FP) detections (number of objects that are falsely detected as nuclei), and the False Negative (FN) detections (number of ground truth nuclei that are not detected). We use the F_1_ score 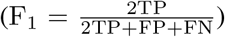 for object-based evaluation. In addition, we adopt the widely used Average Precision (AP), known as the PR curve [67], and the mean Average Precision (mAP) metrics used by the VOC PASCAL [68] and MSCOCO evaluation benchmarks [64].

To reduce evaluation bias given the variability of the segmentation results for different testing data, we use a moving average Intersection-over-Union (IoU) threshold ranging from 0.25 to 0.45 in 0.05 increments (*t* ∈ *T*_IoUs_ = {0.25, 0.3, …, 0.45}). To this end, we define AP_*t*_ and F_1_(*t*) as the AP and F_1_ score evaluated under IoU threshold *t*, respectively. We then define 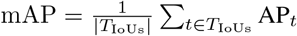 and 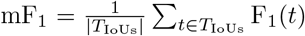 as the mean AP and F_1_ score among all the IoU thresholds, respectively. The evaluation results of our proposed technique and other compared methods on the actual datasets 𝒟_1_ − 𝒟_3_ are given in Table 2. Figure 7 shows the AP of EMR-CNN using BS and AHC (see Section 3.4) with a different number of detectors in an ensemble. All segmentation results and evaluation criteria were verified by a biologist.

**Table 2.** The object-based evaluation results using Average Precision (AP), Mean Average Precision (mAP), and mean F_1_ scores (mF_1_)

| Methods | Microscopy $\mathcal{D}_1$ | | | | Microscopy $\mathcal{D}_2$ | | | | Microscopy $\mathcal{D}_3$ | | | |
| --- | --- | --- | --- | --- | --- | --- | --- | --- | --- | --- | --- | --- |
|  | AP<sub>.25</sub> | AP<sub>.45</sub> | mAP | mF<sub>1</sub> | AP<sub>.25</sub> | AP<sub>.45</sub> | mAP | mF<sub>1</sub> | AP<sub>.25</sub> | AP<sub>.45</sub> | mAP | mF<sub>1</sub> |
| 3D Watershed | 61.51 | 35.12 | 48.87 | 68.31 | 67.58 | 51.20 | 60.30 | 76.36 | 22.33 | 4.62 | 11.96 | 30.22 |
| CellProfiler [25] | 50.46 | 27.35 | 37.79 | 59.41 | 64.26 | 48.66 | 57.52 | 75.09 | 20.46 | 2.85 | 10.15 | 27.75 |
| 3D Squassh [20] | 12.39 | 6.19 | 9.21 | 23.51 | 66.79 | 53.76 | 61.61 | 78.15 | 0.59 | 0.36 | 0.49 | 4.73 |
| VTEA [17] | 45.52 | 38.25 | 42.71 | 61.64 | 64.69 | 45.90 | 57.76 | 74.11 | 25.91 | 10.44 | 17.89 | 38.45 |
| VNet [65] | 77.31 | 61.78 | 70.29 | 82.41 | 53.67 | 34.97 | 45.88 | 64.61 | 37.03 | 12.58 | 23.74 | 41.73 |
| 3D U-Net [66] | 75.56 | 62.97 | 69.63 | 81.81 | 66.89 | 48.14 | 58.12 | 74.62 | 36.89 | 12.40 | 25.19 | 43.39 |
| DeepSynth [30] | 81.57 | 75.59 | 79.51 | 87.19 | 76.42 | 51.86 | 66.19 | 79.42 | 34.98 | 10.91 | 22.83 | 41.77 |
| Cellpose [45] | 81.70 | 81.70 | 81.70 | 89.47 | 71.92 | 64.61 | 68.92 | 80.74 | 45.29 | 17.01 | 30.43 | 51.41 |
| EMR-CNN+BS, M=8 | 82.31 | 69.60 | 76.64 | 86.44 | 77.81 | 70.31 | 75.50 | 85.68 | 53.18 | 33.93 | 45.53 | 64.90 |
| EMR-CNN+AHC, M=8 | **93.19** | **89.46** | **91.04** | **94.95** | **82.37** | **75.71** | **80.13** | **88.15** | **68.26** | **47.65** | **59.61** | **71.05** |

**Figure 7.**
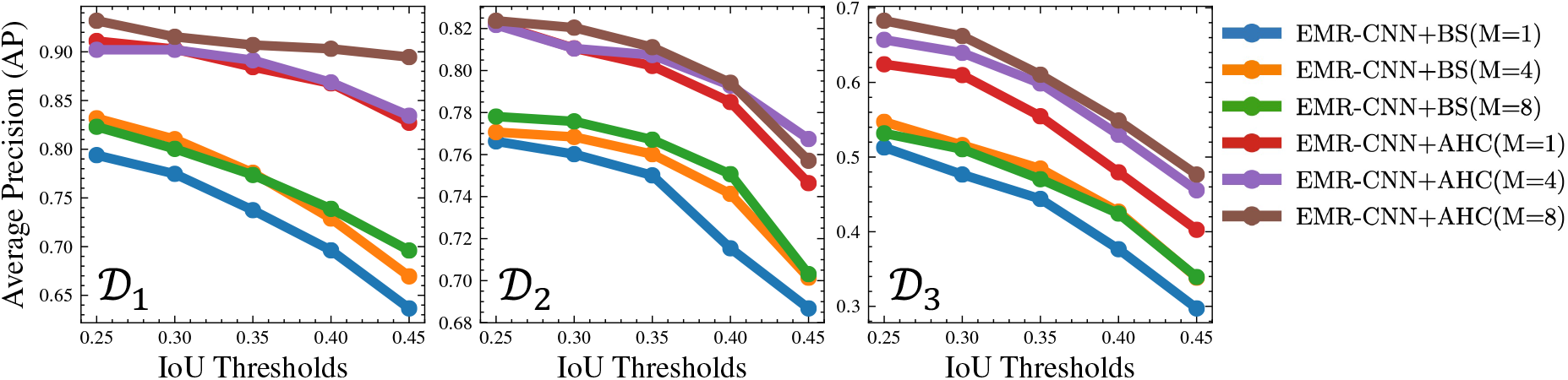
Average Precision (AP) under different IoU thresholds for microscopy 𝒟_1_ − 𝒟_3_. BS and AHC specifies the blob slice or agglomerative hierarchical clustering used for slice fusion module. *M* represents number of detectors in an ensemble.

**Figure 8.**
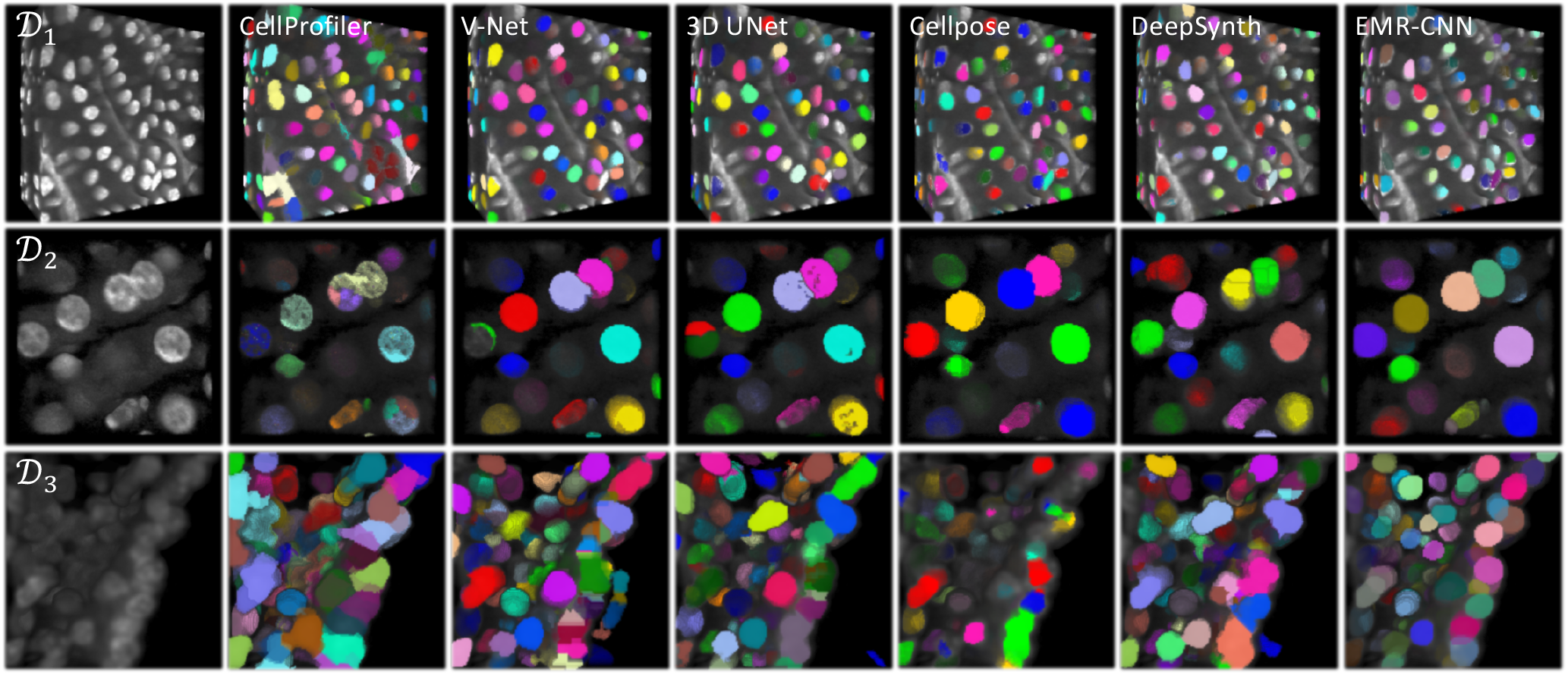
Example testing sub-volumes from original microscopy images and the nuclei instance segmentation results for compared methods. The last column is the segmentation results of our EMR-CNN + AHC, M=8

### 4.3. Running Time Analysis

The running time of our approach and compared methods are given in Table 3. The times reported are based on the training and testing of 𝒟_2_. The compared deep learning methods were trained for 200 epochs on 4 TITAN XP GPUs using the same synthetic data for training EMR-CNN. Our approach takes significantly less training time because each Mask R-CNN model is trained with a subset of the entire training set for only 2000 iterations (2 images/iteration). EMR-CNN and Cellpose take longer time in inferencing since they have to run multiple batches for one volume. The processing of EMR-CNN includes ensemble 2D fusion and 2D to 3D slice fusion. The processing for VNet, UNet, and DeepSynth includes 3D watershed and morphological operations to split touching nuclei.

**Table 3.** Running time analysis. Training: total training time (hours), Inference: model inference time (seconds/volume), Processing: Pre-and Post-processing time (seconds/volume).

| Methods | Microscopy $\mathcal{D}_2$ | | |
| --- | --- | --- | --- |
|  | Training | Inference | Processing |
| CellProfiler | - | - | 30.00 s |
| Squassh | - | - | 30.00 s |
| 3D Watershed | - | - | 4.90 s |
| VTEA | - | - | 2.00 s |
| VNet | 3.07 h | 0.24 s | 3.11 s |
| UNet | 5.13 h | 0.29 s | 3.24 s |
| DeepSynth | 2.53 h | 0.19 s | 3.33 s |
| Cellpose | 4.63 h | 4.48 s | 0.47 s |
| EMR-CNN+BS, M=8 | 0.20 h | 1.01 s | 1.38 s |
| EMR-CNN+AHC, M=8 | 0.20 h | 1.01 s | 1.68 s |

### 4.4. Robustness Analysis

Due to the randomly initialized networks and randomly sampled training set, each detector in the ensemble may produce different False Positive segmentation results for a given image slice. In order to test the stability and robustness of our system, we conducted two experiments. In the first experiment, we randomly add to each detection 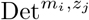, a different number *N* (ranging from 1 to 7) of false positive segmentations of a round shape mask with radius *R* = 4 having a random confidence score between 0.7 and 1.0 (see Figure 9 (left)). In the second experiment, we randomly add *N* = 2 false positive segmentations with different radii *R* ranging from 2 to 8 (see Figure 9 (right)). As the figure indicates, the false positive outliers were removed by our weighted 2D mask fusion with only small losses incurred in the accuracy metrics.

**Figure 9.**
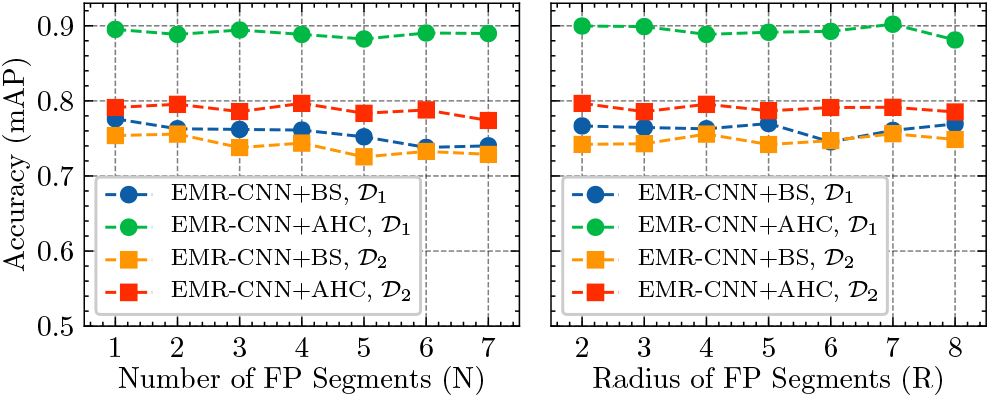
Robustness analysis of proposed method by randomly adding false positive segmentations to each detected image slice

### 4.5. Discussion

Our proposed method was compared with both traditional methods and deep learning-based methods. The traditional methods such as 3D Watershed, CellProfiler, 3D Squash, and VTEA require tuning of many parameters and may work well with one dataset but fail for others. Deep learning-based techniques such as 3D U-Net and DeepSynth are all 3D CNN-based models that directly work on 3D data, whereas Cellpose uses a 2D to 3D reconstruction. In contrast, for the proposed EMR-CNN technique, which is based on Mask R-CNN [54], we utilized two different 2D to 3D slice fusion approaches: Blob-Slice (BS) [57, 17] and Agglomerative Hierarchical Clustering (AHC) [23], with *M* = 1, 4, 8 detectors in an ensemble. As shown in Table 2, the proposed EMR-CNN + AHC approach outperforms the other techniques, based on the AP, mAP, and mF_1_ metrics. As expected, the ensemble with *M* = 8 detectors achieves higher accuracy than *M* = 4 and *M* = 1 for both BS and AHC strategies.

## 5. CONCLUSION

In this paper, we described an ensemble learning and slice fusion strategy for 3D nuclei segmentation. The proposed method uses a weighted 2D mask fusion technique to fuse 2D segmentation masks from different object detectors as well as unsupervised clustering for combining 2D segmentation masks into a 3D segmentation mask. The evaluation results indicate that our approach is stable and robust in the presence of false detections or outliers as well as outperforms some recent 3D CNN-based methods. Moreover, our method does not need any ground truth annotations for training and can inference on any large size volumes. Code has been made available at: http://skynet.ecn.purdue.edu/∼micro/emrcnn/emrcnn_release.zip

## 6. Acknowledgments

This work was partially supported by a George M. OBrien Award from the National Institutes of Health under grant NIH/NIDDK P30 DK079312 and the endowment of the Charles William Harrison Distinguished Professorship at Purdue University. The authors have no conflicts of interest.

